# A rapid method to quantify small-scale vegetation patch structure to complement conventional quadrat surveys

**DOI:** 10.1101/2021.05.12.443830

**Authors:** Liam Butler, Roy A. Sanderson

## Abstract

**Aims:** Vegetation sampling typically involves the use of quadrats, often 1m^2^, to estimate species cover-abundance. Such surveys do not generally record small-scale vegetation patch structure at sub-quadrat scales, for example 10 cm^2^. Here we test a simple method to quantify patch structure that complements conventional techniques. We compare the two methods, and analyse metrics derived from small-scale patch surveys with environment / management data.

**Location:** Northumberland, United Kingdom

**Methods:** We recorded cover-abundance of all species in an upland moor with 1m^2^ quadrats. These were divided into 100 ‘sub-quadrats’, 10 × 10 cm, and the dominant and subdominant species identified. Patch metrics (number, area and shape) for individual species recorded as dominant or subdominant in the sub-quadrat survey were analysed using multivariate generalised linear models with environmental and management data. Sub-quadrat data were also aggregated for each quadrat, to create species composition data. The two sets of compositional data, from whole-quadrat and sub-quadrat aggregations, were compared via Procrustes rotation of ordination scores.

**Results:** Patch number, area and shape for dominant and subdominant species were all significantly affected by soil pH, soil water content, slope and elevation. Effects of proximity to sheep tracks and drainage ditches were less consistent amongst species. Ordinations of vegetation data from conventional and sub-quadrats were similar, with significant Procrustes R-squared of 67% and 70% for dominant and subdominant species respectively.

**Conclusions:** Sub-quadrat surveys can easily be used to complement existing whole-quadrat surveys at little cost in time or resources. Their patch metrics can provide additional insights into the environmental and management drivers that may affect the growth of individual plants or clumps, potentially in relation to plant traits, and thus alter the overall community composition. The methods we describe can readily be adapted to other sizes of quadrats and sub-quadrats in a wide range of vegetation communities.

## 1. Introduction

Vegetation can be measured at multiple scales, for example landscapes with remote-sensed imagery, fields with ground-based habitat surveys, through to conventional quadrats, typically 1 or 2 m. However, individual plants grow and compete with each other at much smaller spatial scales, above- and below-ground, which are more difficult to quantify in the field. Data collected at sub-metre or sub-quadrat spatial scale can provide additional insights into factors that affect patches of each species (Chaneton and Facelli, 1991), which are not visible at coarser spatial scales. Some species can be considered as growing primarily in clumps or patches that occur in a matrix of other species of plants that occur at lower densities and do not necessarily form distinct patches (Fischer and Lindenmayer, 2007; Lejeune et al., 2002). The overall observed patch structure is likely to arise from interactions between these different groups of species and, for example, the physical environment, herbivore grazing, and feedback loops between them.

Patch metrics can be used to assess changes in vegetation patterns at any spatial scale (metres, hectares, or kilometres) and are known to be affected by resource availability, herbivore grazing and species competition (Ritchie, 2010). At field and landscape scales, semi-natural vegetation often creates a patchy discontinuous structure, for example as a result of water availability (Aguiar and Sala, 1999) and topology (Klausmeier, 1999; von Hardenberg et al., 2001). At landscape scales, distinct vegetation patterns have sometimes been reported, with various terminologies such as ‘mosaics’, ‘stripes’, ‘bands’, ‘spots’ or ‘clumps’ (Aguiar and Sala, 1999; Klausmeier, 1999; von Hardenberg *et al*., 2001). One challenge is to develop methods that can encompass a range of spatial scales. For example, Kenkel and Podani, (1991) have highlighted the importance of selecting an appropriate sample size in order to capture spatial pattern; whilst ideally this is larger than the patch size of interest, practical considerations may limit what can be done. In a comprehensive review of the subject, Dale (2000) provides numerous examples of survey methods that have been used at large spatial scales (e.g. field or landscape with quadrats or remote-sensed data), and at small scales (e.g. using point sampling methods).

It is however a challenge to collect multi-species (community) vegetation data across relatively large areas whilst also recording information small-scale spatial structure. Quadrats arranged in a contiguous transect or grid provide comprehensive coverage of the vegetation, for example Bouxin and Gautier (1982) used sixteen 1 m^2^ quadrats placed side-by-side to form a transect in grassland vegetation, where 1 m^2^ was larger than the size of most individual plants. Methods using contiguous quadrats become problematic when surveying large areas, therefore it is not uncommon to have gaps between individual quadrats. Of course, selection of appropriate gap size between quadrats is essential because too large an interval may result in some vegetation mosaics or patches being unsampled (Dale, 2000, p. 36). A popular solution is to arrange non-contiguous quadrats along parallel transects or in a grid (Legendre and Fortin, 1989) which makes the resulting data amenable to a wide range of spatial pattern and interpolation methods for subsequent analysis (Legendre and Legendre, 2012).

Whilst these methods can provide invaluable insights into spatial patterns across relatively large areas of vegetation, they do not provide data on small-spatial patterns within each quadrat. In general, the vegetation quadrat is the unit of sampling, typically via visual estimation of cover of all species that occur within a quadrat (e.g. 1 m^2^) on a quantitative (percentage) or semi-quantitative (Domin) measure. There is evidence that both plant and some animal communities may have patterns at multiple spatial scales, e.g. Carter and O’Connor (1991) report vegetation patterns at both 4 m and 9 m in Savanna grasslands. Quadrat surveys on grids or transects can detect patterns at multiple scales above the size of an individual quadrat using a range of analytical methods, including multi-scale ordination (MSO) (Wagner, 2003) and distance-based Moran’s Eigenvector Maps (MEM) (Dray et al., 2006). However, all these approaches can only detect spatial patterns above the minimum “grain-size” (*sensu* Legendre and Legendre, 2012) that is determined by the actual size of the quadrat *per se*.

Here we describe a simple, rapid approach that can be used in combination with conventional percentage cover quadrat surveys to obtain “within quadrat” information about small-scale spatial patterns, in this case 10 cm. As the scale reduces the number of species of plant within each sample unit declines, so that at small spatial scale in some habitats (e.g. 10 cm cells in upland moors described here) there may only be two to five species present. Thus, visual estimates of percentage cover within a sub-quadrat cell become less meaningful and instead identities of ‘dominant’ and ‘subdominant’ species are likely to be sufficient, whilst still allowing measurement of vegetation patch structure at small scales. One obvious limitation is that it does not detect rare species in, for example a 10 cm cell that would have been recorded by conventional percentage cover abundance at 1m scale. Thus, sub-quadrat surveys should be viewed as complementary to conventional percentage cover surveys.

The overall aim of this research was to test the utility of small-scale survey methods as a means to understand patch-level community composition. Specific objectives were to:

a. Field-test a method for within-quadrat surveys, and calculate small-scale measures of vegetation patch number, area and shape and quantify their relationship to local environmental and management conditions.
b. Compare species assemblage data from conventional percentage-cover whole-quadrat (1m-scale) surveys to assemblages derived from within-quadrat (10 cm-scale) assessments of the incidence of the most common species.

## 2. Materials and Methods

### 2.1 Study site

Vegetation and environmental data were collected from a 96 ha area of sheep-grazed semi-natural grassland and moor (known as ‘Ashtrees Dipper’) at Cleughbrae Farm, Northumberland, UK (NY 83301 96079; Fig. 1). Most of the vegetation is in UK National Vegetation Classification (NVC) classes *Calluna vulgaris*-*Deschampsia flexuosa* heath (H9), *Scripus cespitosus*-*Erica tetralix* wet heath (M15), *Deschampsia flexuosa* grassland (U2), *Festuca ovina-Agrostis capillaris-Galium saxatile* grassland (U4) and *Nardus stricta-Galium saxatile* grassland (U5) habitats (Rodwell, 1992; Rushton et al., 1992; Smith et al., 1992; Sanderson et al., 1995). It is north-facing, altitudes 250 m to 350 m and is characterised by acidic soils, primarily surface-water gleys, podzols and raw peats (Cranfield University, 2017) with high overall water content.

**Fig. 1.**
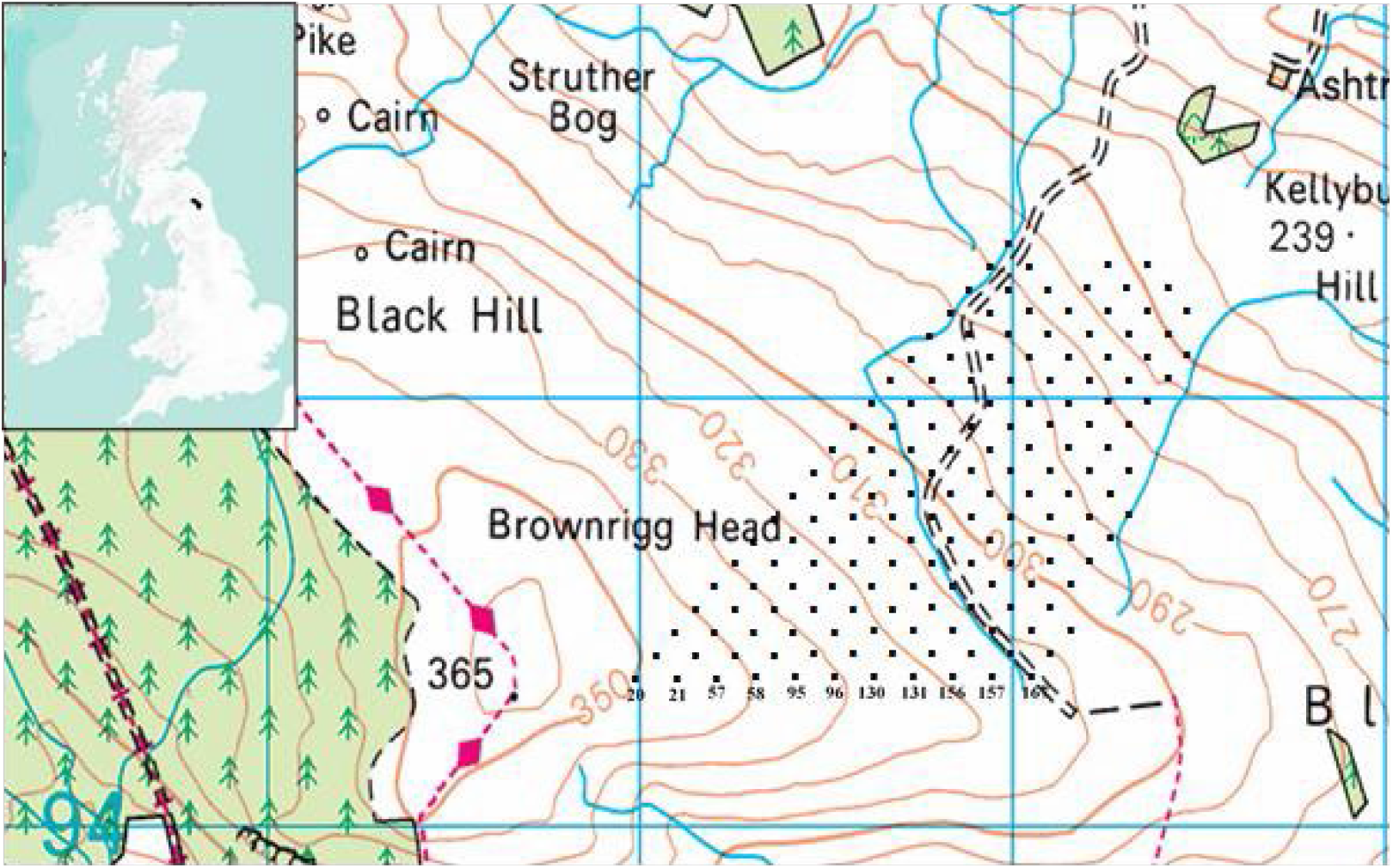
Location of the study site (Ashtrees) in the NE UK. The top left image shows the location of Catchment of River Rede. The points show the location of the 167 quadrats along transects. Quadrats were placed 75m apart. The squares are 1km Ordnance Survey.

### 2.2 Collection and management of vegetation data at 10cm scale

Vegetation was surveyed from 167 1m^2^ quadrats placed along transects 150m apart, with quadrats 75m apart along each transect (Fig. 1). Each quadrat was sub-divided into 100 cells 10 cm × 10 cm via wires stretched across the frame, henceforth referred to as a ‘sub-quadrat’. Vegetation cover was recorded as visual estimates of percentage cover of the whole quadrat, for all species, which is a widely used conventional assessment. In addition, within each sub-quadrat the incidence of the most abundant (‘dominant’) and second most abundant (‘subdominant’) species of vegetation were recorded, along with their spatial locations in each of the 100 sub-quadrats. All surveys were completed in 1991.

The final step was to identify different vegetation patches for each species within a quadrat, encoding each patch separately. Diagonally adjoining cells were assumed to belong to the same patch. An example of the output for a single species in a quadrat containing five patches is provided in Fig. 2.

**Fig. 2.**
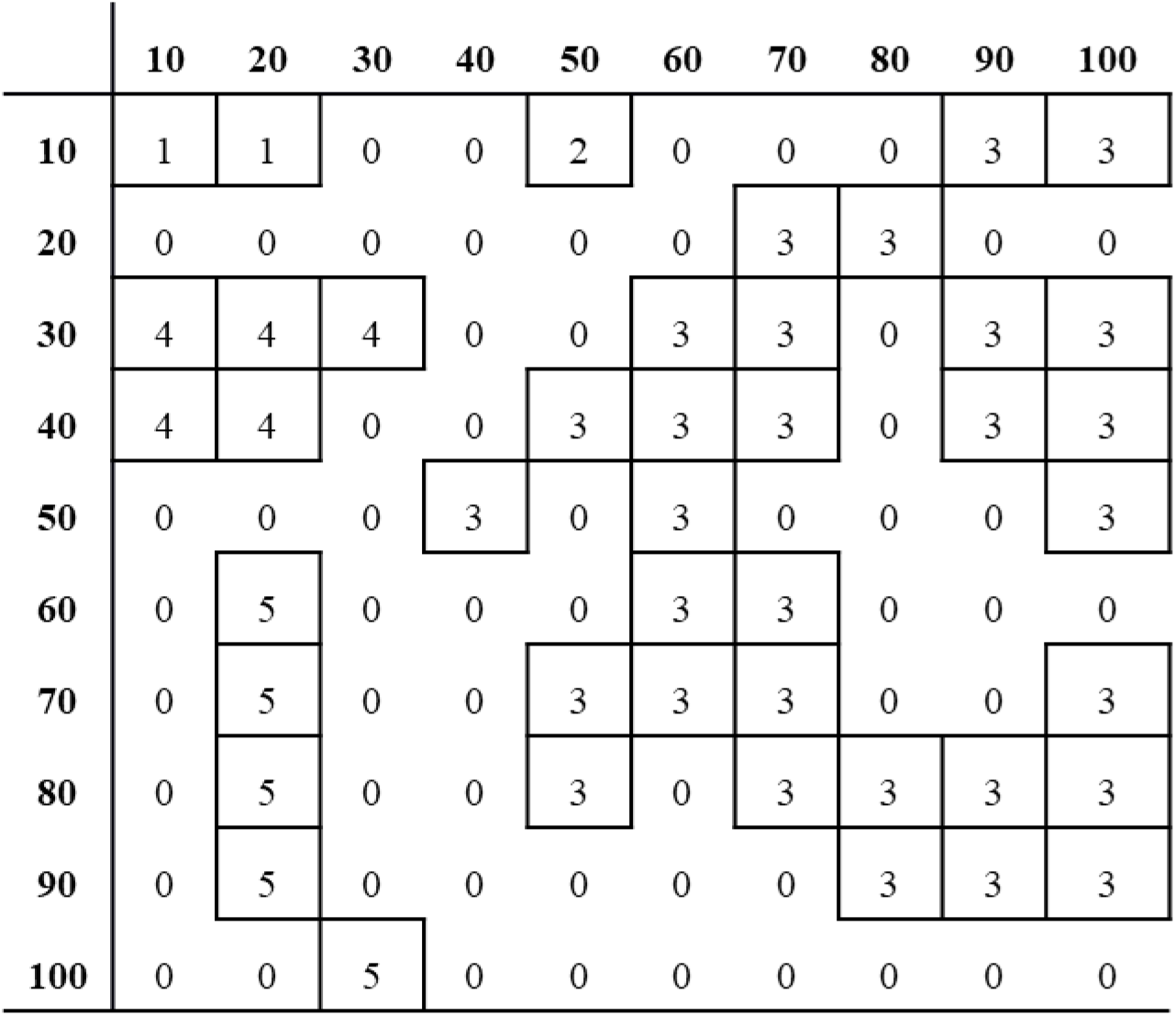
Matrix of patch numbers for a single species in a quadrat. Each number (1 - 5) is a different patch formed by the species and 0 is background of other species.

### 2.1 Vegetation patch metrics

Large numbers of metrics that differ in complexity have been developed to quantify different aspects of spatial patterns, in particular those popularised by the FRAGSTATS software by McGarigal and Marks (1994) for landscape-scale spatial analysis. No single metric can describe all aspects of spatial structure, and we have chosen to focus on three metrics that are widely used and relatively simple to interpret: the number of vegetation patches, the mean patch area, and the shape index. All three were calculated separately for each dominant or subdominant species surveyed (see Supplementary Material for examples), but for simplicity we only report overall measures for all species in each 1m^2^ quadrat. The shape index (Equation 1) provides an alternative to fractal dimension (Ritchie, 2009; McGarigal, 2017) and measures the complexity of a two-dimensional object compared to a standard shape, generally a circle (for vector data) or square (for raster data, as collected in this study).

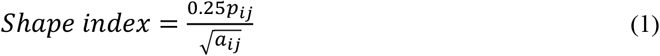

where

*P_ij_* = perimeter of patch *ij*
*a_ij_* = area of patch *ij*

This has been widely used in landscape ecology (Forman and Godron, 1981); note that it is not size-dependent, and the index equals 1 for square patches of any size and increases without limit as the patch becomes increasingly non-square (i.e., more geometrically complex; Fig. 3; McGarigal, 2017).

**Fig. 3.**
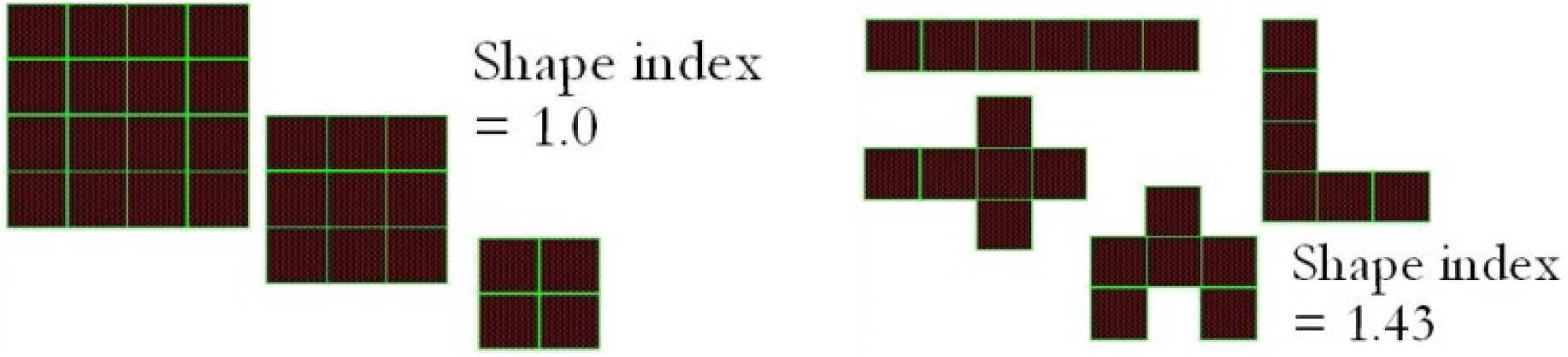
Example and explanation of shape index where 1 = square of any size. Adopted from McGarigal, 2017.

### 2.2 Variation in patch metrics in response to environment and management

Multivariate generalised linear models (Warton et al., 2012; Wang et al., 2017) were used to measure the relationships between the three vegetation patch metrics with environmental and management conditions. Multivariate GLMs were undertaken with the ‘mvabund’ package (Wang et al., 2017) with Gaussian error models; the technique provides information for both individual species, as well as the collective response of all species to the predictors. Environmental predictor variables were soil pH, soil water content, altitude and slope (see Sanderson et al 1995 for details). Positions of sheep paths and drainage grips were digitised in ESRI ArcGIS 10.6 from stereo-pairs of aerial photographs taken in 1991 that encompassed the whole study site. The distance to nearest sheep track and drainage ditch were determined in ArcGIS; circular buffers of 10m, 25m and 35m radius around each quadrat were created and the total length of sheep track in each buffer calculated. These variables (distance to nearest ditch, distance to nearest sheep track, and length of sheep track within the buffers) were used as additional predictor variables in the multivariate GLMs, to determine effects of drainage management and grazing livestock activity patterns across the site. Multivariate GLM was undertaken on patch metrics derived from all the dominant and subdominant species (see Supplementary Information) but are only reported in detail for the eleven most common species: *Calluna vulgaris*, *Eriophorum vaginatum*, *Juncus effusus*, *Juncus squarrosus*, *Molinia caerulea*, *Nardus stricta*, *Carex nigra*, *Deschampsia flexuosa*, *Galium saxatile*, *Potentilla erecta* and *Vaccinium myrtillus*. Some of these species, e.g. *C. vulgaris, E. vaginatum, J. effusus, J. squarrosus, M. caerulea* and *N. stricta*, typically form distinct clumps, whilst others e.g. *C. nigra, D. flexuosa, G. saxatile, P. erecta, V. myrtillus*, often form a background vegetation matrix (Grime, 1978; Robinson and Rorison, 1983; Rushton *et al*., 1992; Smith *et al*., 1992; Britton et al., 2003).

### 2.3 Comparison of sub-quadrat data with conventional survey methods

One risk with the sub-quadrat survey method is that it is less likely to detect rare spaces with low cover-abundance. These rare species will be recorded in a conventional percentage cover survey of the whole quadrat, but may be omitted entirely from a within-quadrat survey of the same quadrat, unless they happen to be particularly abundant in one 10 cm. Therefore, whilst the sub-quadrat method provides a means to assess small-scale spatial pattern, it is important to determine whether the methodology is representative of differences in vegetation composition across the site as a whole. The latter is more accurately measured via the conventional percentage cover survey in this case study.

The numbers of 10 cm cells occupied by each species for either the dominant or the subdominant species were totalled, such the sum across all species was a value between 0 and 100. This produced a measure of the overall ‘incidence’ of each species, as either a dominant or subdominant. In contrast, conventional percentage cover captured all species (including relatively rare ones with low cover abundance), and the overall cover abundance was sometimes greater than 100% due to overlapping layers of vegetation. Non-metric multidimensional scaling (NMDS) was used to summarise vegetation assemblages derived from the dominant or subdominant surveys. These two NMDS ordinations of the dominant and subdominant species were compared with the equivalent ordination of raw percentage cover obtained from the conventional whole-quadrat survey using Procrustes rotation (Gower, 1975). Procrustes rotation rescales and rotates ordinations so that they match as closely as possible (Gower, 1975), and can be used to test the similarity m^2^ (Procrustes residual derived from the sum of the squared deviation) and R^2^ (correlation coefficient) of each pair of ordinations. Two comparisons were made: i) species cover derived from percentage visual estimates versus dominant vegetation (0-100) and ii) species cover derived from percentage visual estimates versus subdominant vegetation (0-100), using the R ‘vegan’ package (Oksanen et al., 2015).

## 3. Results

Overall analysis of patch metrics for all species that occurred as dominant and subdominants indicated strong associations with the environment (Table 1). The number of patches was greater on neutral soils, and steeper slopes, and smaller at lower altitude, for both dominants and subdominant species. Patch area was larger on acid soils, whilst patch shape complexity was higher on wet soils at higher altitude. The presence of sheep tracks within 10 m of a quadrat resulted in a greater number of vegetation patches. Overall patterns were broadly similar for species when recorded at both dominant and subdominants, although exceptions included soil pH and shape complexity, and soil water content with the number of vegetation patches.

**Table 1.** Summary of overall multivariate GLM analysis for dominant and subdominant vegetation with Cohen’s *d* effect size and *p* significance of environmental variables. ST = Sheep Tracks.

| Env. Variable | Dominant |  |  |  |  |  | Subdominant |  |  |  |  |  |
| --- | --- | --- | --- | --- | --- | --- | --- | --- | --- | --- | --- | --- |
|  | Number |  | Area |  | Shape |  | Number |  | Area |  | Shape |  |
| | $d$ | $p$ | $d$ | $p$ | $d$ | $p$ | $d$ | $p$ | $d$ | $p$ | $d$ | $p$ |
| Soil pH | 1.434 | **0.001** | -2.288 | **0.001** | 0.120 | **0.001** | 0.313 | **0.001** | -0.667 | **0.001** | -0.024 | **0.001** |
| Slope | 0.746 | **0.001** | 0.063 | 0.069 | -0.027 | **0.001** | 0.506 | **0.019** | -0.398 | **0.036** | -0.075 | **0.001** |
| % soil water content | 0.364 | **0.001** | -0.010 | **0.029** | 0.124 | **0.001** | -0.497 | **0.002** | 0.134 | **0.033** | 0.189 | **0.001** |
| Altitude | -0.564 | **0.001** | 0.051 | **0.053** | -0.220 | **0.048** | -0.270 | **0.011** | 0.394 | **0.066** | 0.317 | **0.001** |
| ST length (10m) | 0.002 | 0.720 | -0.133 | 0.430 | 0.003 | 0.533 | 0.008 | 0.436 | 0.027 | 0.362 | 0.127 | 0.362 |
| ST distance (10m) | 0.548 | **0.024** | -0.193 | 0.085 | -0.030 | 0.478 | 0.017 | **0.048** | -0.140 | 0.052 | -0.025 | 0.052 |
| ST length (25m) | -0.002 | 0.060 | 0.034 | 0.544 | -0.603 | 0.055 | -0.237 | 0.281 | 0.195 | 0.346 | 0.231 | 0.346 |
| ST distance (25m) | -0.012 | 0.110 | -0.231 | 0.875 | -0.068 | 0.289 | 0.007 | 0.466 | 0.018 | 0.400 | 0.192 | 0.420 |
| ST length (35m) | 0.012 | 0.200 | -0.183 | 0.169 | -0.061 | 0.097 | 0.349 | 0.206 | -0.355 | 0.137 | -0.18 | 0.085 |
| ST distance (35m) | 0.066 | 0.071 | 0.231 | **0.023** | 0.603 | 0.059 | 0.006 | 0.205 | -0.019 | 0.196 | -0.192 | 0.202 |
| Ditch distance | -0.208 | 0.066 | 0.149 | 0.167 | -0.119 | **0.023** | 0.306 | 0.063 | 0.232 | 0.108 | 0.056 | 0.086 |

Multivariate GLM of the eleven most common species showed relatively few strong associations with these predictors (see Supplementary Tables S1 for full breakdown of results for each species). *E. vaginatum* was affected most, with its number of patches being negatively related to slope, and positively related to soil water content. *E. vaginatum* patch area and shape index increased with soil water content, but shapes were less complex on steeply sloping land. *J. effusus* had less complex patches (as measured by shape index) at higher altitude. Patch areas of *D. flexuosa* increased with soil water content, and both the number and areas of patches were higher on more acidic soils. There was evidence that proximity of sheep tracks, especially within 10m of a quadrat, affected the patch structure, and (weaker) evidence that distance to the nearest drainage ditch also had an effect.

A total of 58 species were recorded across all 167 quadrats by conventional whole-quadrat 1m^2^ assessments, compared to only 41 and 51 species for the dominant and subdominant species respectively at the 10 cm sub-quadrat scale. Nevertheless, comparison of the overall community composition based on Procrustes analysis of the NMDS scores showed strong agreement between all survey methods (Table 2). All three ordination plots (Supplementary Figure S1) displayed similar patterns, with *Calluna vulgaris*, *Eriophorum vaginatum* and *Sphagnum* species being at the extremes of NMDS Axis 1 and *Holcus mollis*, *Juncus articulatus* and *Cardemine pratensis* at the extremes of Axis 2.

**Table 2.** Summary of Procrustes rotation comparison between i) percentage cover data, denoted as ‘survey’, and dominant patch area data, ii) survey versus subdominant vegetation patch area data and iii) dominant vs subdominant vegetation patch area data. ‘m^2^’ is Procrustes residual, R = Procrustes correlation coefficient.

| Procrustes rotation comparison | m<sup>2</sup> | R | p-value |
| --- | --- | --- | --- |
| Survey vs Dominant | 0.5565 | 0.6659 | 0.001 |
| Survey vs Subdominant | 0.5111 | 0.6992 | 0.001 |
| Dominant vs Subdominant | 0.5255 | 0.6888 | 0.001 |

## 4. Discussion

Patch metrics have been widely used to quantify spatial structure in habitats at landscape scales, but their implementation across large areas at small spatial scales has been much rarer. The concept of ‘micropatterning’ of vegetation was introduced in the mid-1980s (Ohsawa, 1984) but the method did not compare vegetation patterns within different environments or between vegetation species (Ohsawa, 1984). Berg *et al*. (1997) measured the effects of environment and management on vegetation patch fragmentation at field scale in sheep-dominated areas. Here, we demonstrate the utility of using patch metrics at sub-metre scales, via vegetation surveys that employed a simple, rapid, inexpensive field method that can also be used to derive multi-species assemblages for conventional community-level analyses. In the field the within-quadrat survey method proved to be extremely fast, especially as it was done immediately after the conventional survey of each quadrat. In practice, recording the additional information at 10 cm scale took an additional 4 to 5 minutes of survey time per quadrat.

One challenge to undertaking within-quadrat studies is that as the spatial scale of the survey unit becomes smaller the total number of species surveyed is also reduced. Thus, our dominant and subdominant assessments recorded 29% and 12% fewer species respectively than the traditional whole-quadrat approach. This drop in species richness might suggest that the within-quadrat method does not provide reliable data when comparing spatial patterns within different quadrats. Nevertheless, the high correlation of the community summaries (from NMDS and Procrustes) indicates that the within-quadrat survey method provided a comparable method to determine similarity of community composition between quadrats. This accords with Kokvidze and Ohsawa (2002) who suggested that community-level analyses were particularly affected by the most frequent species. We found strong similarities in the results for both the dominant and sub-dominant data, and so it is possible that the latter could be omitted (e.g. to reduce field survey times) without compromising the quality of the data. Pilot studies would, however, be needed in other habitats before this could be recommended in general. It is important to stress that the traditional whole-quadrat and sub-quadrat methods should be considered as complementary, rather than alternatives to each other. Conventional whole quadrat surveys are essential for more accurate species inventories. If one aim of a survey is to allocate vegetation to an extant nationwide community classification system, such as the UK National Vegetation Classification (Rodwell, 1992), then whole-quadrat surveys are essential.

The number of patches formed was greater for the dominant than subdominant species. To some extent this is to be expected, as species that are dominant in an area tend to form larger patches, occupying more space, and be less fragmented compared to subdominants. Whilst some species in a quadrat were mainly recorded as dominants (e.g. *Hypnum cupressiforme*), or as sub-dominants (e.g. *Lolium perenne, Poa trivialis*), others could be either dominant or subdominant, depending on their location within an individual quadrat (e.g. *Molinia caerulea, Carex nigra*). Those species that were primarily dominant in a quadrat were more likely to form large clumps whilst those primarily sub-dominant were more likely to form fragmented matrices or tufts. Variation in patch structure might arise from species phenology, functional traits, competitive abilities, regenerative strategies and seed dispersal (Grime et al., 1988). These differences in expansion and fragmentation depend on both the characteristics of the individual plant species, and the environment in which it grows. For example, root competition within and between species may also play a role in patch formation and development, although of course these can be difficult to quantify (Herben et al., 2020).

Multivariate GLM analysis of both dominant and subdominant species indicated significant relationships between the number of patches and the environment, especially soil pH and soil water content. The area of dominant patches increased with altitude, whilst the shapes of subdominant patches became more complex (Table 2); to some extent this may reflect competition between the dominant and subdominant species occupying the same space (Addicott et al., 1987; Grime, 1988; Suding et al., 2008). Relationships between slope and vegetation are likely to be site-specific and therefore it can be difficult to predict the effects on patch metrics in other habitats (Bennie et al., 2008). Our results indicate that subdominant patches were more affected by slope than dominant species. Furthermore, in both cover types, patch shape became less complex and overall patch area decreased with increased slope, which suggests that species became more fragmented. Acid soils were associated with larger patches of species adapted to these conditions and decreased the number of fragmented patches across the field (Addicott et al., 1987). Soil pH is affected by water content, and as different species are adapted to wet or dry soils, this will partly determine which ones become dominant as a result of interspecific competition and microtopography (Wigmosta et al., 2002).

The number of patches formed by dominant or subdominant vegetation species increased when sheep tracks occurred within 10m of the quadrat, whilst subdominants also had increased shape complexity. These two findings may suggest that high grazing pressure increases vegetation fragmentation, which accords with numerous previous studies (Thomas, 1959; Plumptre, 1994; Maron and Crone, 2006; Stewart and Pullin, 2006). Of course, the vicinity of sheep tracks should not be viewed as a simple surrogate for grazing pressure, as tracks are sometimes merely used for ‘transit’ by animals. However, it was noticeable that the effects of sheep tracks were only apparent at the shortest buffer distance of 10m. Species near sheep tracks might be less palatable to sheep and more resistant to trampling, which might also increase shape complexity of the nearby vegetation (Maron and Crone, 2006). Furthermore, increases in pattern complexity has been reported as an ‘evasion’ strategy to optimise chances of survival for some species in areas of high grazing effort (Fisher *et al*., 1996). This ‘evasion’ has been observed in some of the more palatable species (e.g. *Caluna vulgaris, Nardus stricta*) allowing other species such as *Agrostis capillaris*, *Deschampsia flexuosa*. *Festuca rubra* etc. to grow in proximity to sheep tracks while the palatable species grow further away from the tracks (Fenton, 1937; Krahulec *et al*., 2001)

New drainage ditches are no longer actively dug across large areas of the UK uplands but historical and extant drainage networks still affect vegetation growth (Holden *et al*., 2004; Ramchunder *et al*., 2009). While we did not find consistent associations between proximity to drainage ditches and patch metrics, drainage ditches can still influence the abundance of mosses and unpalatable grasses such as *Nardus stricta* (Coulson *et al*., 1990; Ramchunder *et al*., 2009). The overall number of patches formed by dominant species decreased when in close proximity to a ditch and shape became less complex (Table 2); the mechanisms behind this are unclear, but might arise from lower soil water favouring other species.

## Conclusion

Dominant/subdominant within-quadrat surveys provide a robust yet cost-effective method to quantify vegetation patches structure at small spatial scales. The method does not record rarer species with low cover-abundance, therefore it should always be considered a complementary technique to whole-quadrat surveys, and never a replacement. Nevertheless community-level analyses indicate that it provides robust data to investigate vegetation responses to environmental and management determinants of spatial pattern. The approach we describe can be readily adapted to predict potential patch metrics for individual species, or the vegetation as a whole, e.g. across a whole field via conventional interpolation methods. The time required for a sub-quadrat survey is relatively low, meaning that it can be readily incorporated as an additional tool for practising field ecologists.

## Acknowledgements

The authors thank Prof. Stephen P. Rushton and Dr Gavin Stewart from Newcastle University, and Dr Oliver Pescott from the Centre for Ecology and Hydrology, for advice on this project and analyses.

## Data availability

The data that support the findings of this study are openly available in the Newcastle University Data Repository / FigShare at https://doi.org/10.25405/data.ncl.12623816.

## References

Addicott, J. F., Aho, J. M., Antolin, M. F., Padilla, D. K., Richardson, J. S. and Soluk, D. A. (1987) ‘Ecological neighborhoods: scaling environmental patterns’, Oikos, pp. 340–346.

Aguiar, M. R. and Sala, O. E. (1999) ‘Patch structure, dynamics and implications for the functioning of arid ecosystems’, Trends in Ecology & Evolution, vol. 14, no. 7, pp. 273–277.

Bennie, J., Huntley, B., Wiltshire, A., Hill, M. O. and Baxter, R. (2008) ‘Slope, aspect and climate: Spatially explicit and implicit models of topographic microclimate in chalk grassland’, Ecological Modelling, vol. 216, no. 1, pp. 47–59 [Online]. DOI: 10.1016/j.ecolmodel.2008.04.010.

Berg, G., Esselink, P., Groeneweg, M. and Kiehl, K. (1997) ‘Micropatterns in Festuca rubra-dominated salt-marsh vegetation induced by sheep grazing’, Plant Ecology, vol. 132, no. 1, pp. 1–14.

Bouxin, G. and Gautier, N. (1982) ‘Pattern analysis in Belgian limestone grasslands’, Vegetatio, vol. 49, no. 2, pp. 65–83 [Online]. DOI: 10.1007/BF00052760.

Britton, A., Marrs, R., Pakeman, R. and Carey, P. (2003) ‘The influence of soil-type, drought and nitrogen addition on interactions between Calluna vulgaris and Deschampsia flexuosa: implications for heathland regeneration’, Plant Ecology, vol. 166, no. 1, pp. 93–105.

Carter, A. J. and O’Connor, T. G. (1991) ‘A two-phase mosaic in a savanna grassland’, Journal of Vegetation Science, vol. 2, no. 2, pp. 231–236 [Online]. DOI: https://doi.org/10.2307/3235955.

Chaneton, E. J. and Facelli, J. M. (1991) ‘Disturbance effects on plant community diversity: spatial scales and dominance hierarchies’, Vegetatio, vol. 93, no. 2, pp. 143–155 [Online]. DOI: 10.1007/BF00033208.

Coulson, J. C., Butterfield, J. E. L. and Henderson, E. (1990) ‘The effect of open drainage ditches on the plant and invertebrate communities of moorland and on the decomposition of peat’, Journal of Applied Ecology, pp. 549–561.

Dale, M. R. (2000) Spatial pattern analysis in plant ecology, Cambridge university press.

Dray, S., Legendre, P. and Peres-Neto, P. R. (2006) ‘Spatial modelling: a comprehensive framework for principal coordinate analysis of neighbour matrices (PCNM)’, Ecological Modelling, vol. 196, no. 3, pp. 483–493 [Online]. DOI: 10.1016/j.ecolmodel.2006.02.015.

Fenton, E. W. (1937) ‘The influence of sheep on the vegetation of hill grazings in Scotland’, Journal of Ecology, vol. 25, no. 2, pp. 424–430.

Fischer, J. and Lindenmayer, D. B. (2007) ‘Landscape modification and habitat fragmentation: a synthesis’, Global Ecology and Biogeography, vol. 16, no. 3, pp. 265–280 [Online]. DOI: 10.1111/j.1466-8238.2007.00287.x.

Fisher, A. S., Podniesinski, G. S. and Leopold, D. J. (1996) ‘Effects of drainage ditches on vegetation patterns in Abandoned agricultural peatlands in central New York’, Wetlands, vol. 16, no. 4, pp. 397–409 [Online]. DOI: 10.1007/BF03161329.

Forman, R. T. and Godron, M. (1981) ‘Patches and structural components for a landscape ecology’, BioScience, vol. 31, no. 10, pp. 733–740.

Grime, J. P. (1988) ‘The CSR model of primary plant strategies—origins, implications and tests’, in Plant evolutionary biology, Springer, pp. 371–393.

Grime, J. P., Hodgson, J. G. and Hunt, R. (1988) ‘Comparative Plant Ecology: A Functional Approach to Common British Species.(Agrostis spp., pp. 58–65.) Unwin Hyman’, London, England, UK.

von Hardenberg, J., Meron, E., Shachak, M. and Zarmi, Y. (2001) ‘Diversity of vegetation patterns and desertification’, Physical Review Letters, vol. 87, no. 19, p. 198101.

Herben, T., Balšánková, T., Hadincová, V., Krahulec, F., Pecháčková, S., Skálová, H. and Krak, K. (2020) ‘Fine-scale root community structure in the field: species aggregations change with root density’, Journal of Ecology.

Holden, J., Chapman, P. J. and Labadz, J. C. (2004) ‘Artificial drainage of peatlands: hydrological and hydrochemical process and wetland restoration’, Progress in Physical Geography, vol. 28, no. 1, pp. 95–123 [Online]. DOI: 10.1191/0309133304pp403ra.

Kenkel, N. C. and Podani, J. (1991) ‘Plot size and estimation efficiency in plant community studies’, Journal of vegetation science, Wiley Online Library, vol. 2, no. 4, pp. 539–544.

Klausmeier, C. A. (1999) ‘Regular and irregular patterns in semiarid vegetation’, Science, vol. 284, no.5421, pp. 1826–1828.

Krahulec, F., Skálová, H., Herben, T., Hadincová, V., Wildová, R. and Pecháčková, S. (2001) ‘Vegetation changes following sheep grazing in abandoned mountain meadows’, Applied Vegetation Science, vol. 4, no. 1, pp. 97–102 [Online]. DOI: 10.1111/j.1654-109X.2001.tb00239.x.

Legendre, P. and Fortin, M. J. (1989) ‘Spatial pattern and ecological analysis’, Vegetatio, vol. 80, no. 2, pp. 107–138 [Online]. DOI: 10.1007/BF00048036.

Legendre, P. and Legendre, L. (2012) Numerical Ecology, Elsevier.

Lejeune, O., Tlidi, M. and Couteron, P. (2002) ‘Localized vegetation patches: A self-organized response to resource scarcity’, Physical Review E, American Physical Society, vol. 66, no. 1, p. 010901 [Online]. DOI: 10.1103/PhysRevE.66.010901.

Maron, J. L. and Crone, E. (2006) ‘Herbivory: effects on plant abundance, distribution and population growth’, Proceedings of the Royal Society B: Biological Sciences, vol. 273, no. 1601, pp. 2575–2584 [Online]. DOI: 10.1098/rspb.2006.3587.

McGarigal, K. (2017) ‘Landscape metrics for categorical map patterns’, Lecture Notes. Available online: http://www.umass.edu/landeco/teaching/landscape_ecology/schedule/chapter9_metrics.pdf (accessed on 3 July 2018).

Ohsawa, M. (1984) ‘Differentiation of vegetation zones and species strategies in the subalpine region of Mt. Fuji’, Vegetatio, vol. 57, no. 1, pp. 15–52 [Online]. DOI: 10.1007/BF00031929.

Plumptre, A. J. (1994) ‘The effects of trampling damage by herbivores on the vegetation of the Pare National des Volcans, Rwanda’, African Journal of Ecology, vol. 32, no. 2, pp. 115–129.

Ramchunder, S. J., Brown, L. E. and Holden, J. (2009) ‘Environmental effects of drainage, drain-blocking and prescribed vegetation burning in UK upland peatlands’, Progress in Physical Geography, vol. 33, no. 1, pp. 49–79 [Online]. DOI: 10.1177/0309133309105245.

Ritchie, M. E. (2009) Scale, heterogeneity, and the structure and diversity of ecological communities, Princeton University Press, vol. 47.

Robinson, D. and Rorison, I. H. (1983) ‘A comparison of the responses of Lolium perenne L.*, Holcus lanatus L. and Deschampsia flexuosa (L.) Trin. To a localised supply of nitrogen’, New Phytologist, vol. 94, no. 2, pp. 263–273.

Rushton, S. P., Byrne, J. P. and Young, A. G. (1992) ‘Redesdale: a preliminary test catchment for the NERC/ESRC Land Use Programme’, ITE symposium.

Smith, R. S., Rushton, S. P. and Wadesworth, R. A. (1992) ‘Predicting vegetation change in an upland environment’, ITE symposium.

Stewart, G. B. and Pullin, A. S. (2006) ‘Does sheep-grazing degrade unimproved neutral grasslands managed as pasture in lowland Britain’, Systematic Review, vol. 15.

Suding, K. N., Lavorel, S., Chapin Iii, F. S., Cornelissen, J. H., DIAz, S., Garnier, E., Goldberg, D., Hooper, D. U., Jackson, S. T. and Navas, M.-L. (2008) ‘Scaling environmental change through the community-level: a trait-based response-and-effect framework for plants’, Global Change Biology, vol. 14, no. 5, pp. 1125–1140.

Thomas, A. S. (1959) ‘SHEEP PATHSO: Observations on the variability of chalk pastures’, Grass and Forage Science, vol. 14, no. 3, pp. 157–164.

Wagner, H. H. (2003) ‘Spatial Covariance in Plant Communities: Integrating Ordination, Geostatistics, and Variance Testing’, Ecology, vol. 84, no. 4, pp. 1045–1057 [Online]. DOI: https://doi.org/10.1890/0012-9658(2003)084[1045:SCIPCI]2.0.CO;2.

Wang, X., Gao, Q., Wang, C. and Yu, M. (2017) ‘Spatiotemporal patterns of vegetation phenology change and relationships with climate in the two transects of East China’, Global Ecology and Conservation, vol. 10, pp. 206–219 [Online]. DOI: 10.1016/j.gecco.2017.01.010.

Warton, D. I., Wright, S. T. and Wang, Y. (2012) ‘Distance-based multivariate analyses confound location and dispersion effects’, Methods in Ecology and Evolution, vol. 3, no. 1, pp. 89–101 [Online]. DOI: 10.1111/j.2041-210X.2011.00127.x.

Wigmosta, M. S., Nijssen, B., Storck, P. and Lettenmaier, D. P. (2002) ‘The distributed hydrology soil vegetation model’, Mathematical models of small watershed hydrology and applications, pp. 7–42.

