## Supplementary figures and tables for "A rapid method to quantify small-scale vegetation patch structure to complement conventional quadrat surveys"

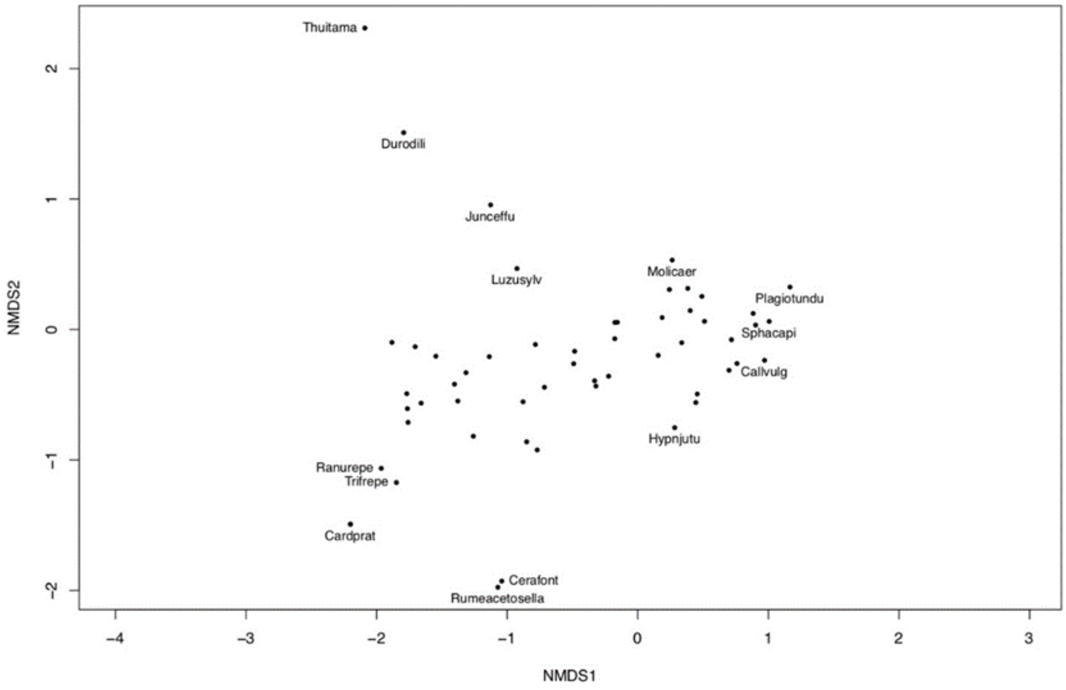


**Fig. S.1a** NMDS ordination species plot from percentage cover abundance data at Ashtrees. Thuitama = *Thuidium tamariscinum*, Durodili = *Dryopteris dilitata*, Juneffu = *Juncus effusus*, Luzusylv = *Luzula sylvatica,* Molicaer = *Molinia caerulea*, Plagiotundu = *Plagiothecuim undulatum*, Sphacapi = *Sphagnum capillofolium,* Callvulg = *Calluna vulgaris*, Hypnjutu = *Hypnum jutlandicum*, Cetafont = *Cerastium fontanum*, Rumeacetosella = *Rumexacetosella*, Cardprat = *Cardemine pratensis*, Trifrepe = *Trifolium repens*, Ranurepe = *Ranunculus repens*

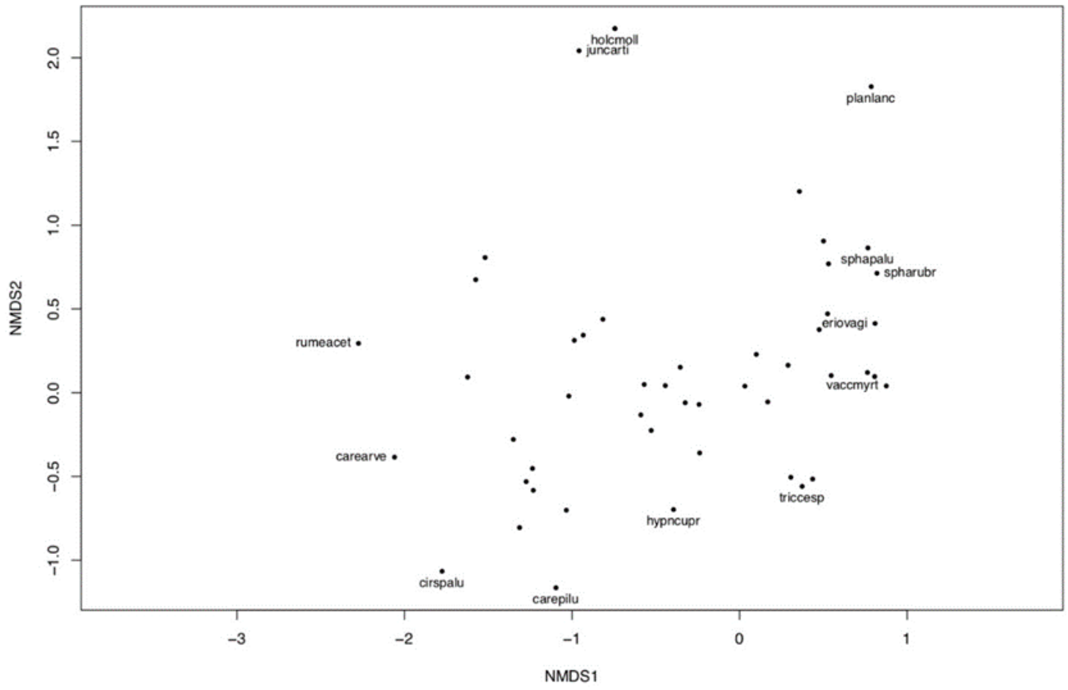


**Fig. S.1b** NMDS ordination species plot for dominant vegetation species found at Ashtrees. Holcmolli = *Holcus lanatus*, planlanc = *Plantego lanceolata*, sphapalu = *Sphagnum palustre*, spharubr = *Sphagnum rubra*, eriovagi = *Eriophorum vaginatum*, vaccmyrt = *Vaccinium myrtillus*, triccesp = *trichophorum cespitosum*, hypncupr = *Hypnum cupressiforme*, carepilu = *Carex pilulifera,* cirspalu = *Cirsium palustre*, Carearve = *Cirsium arvense*, rumeacet = *Rumex acetosa*

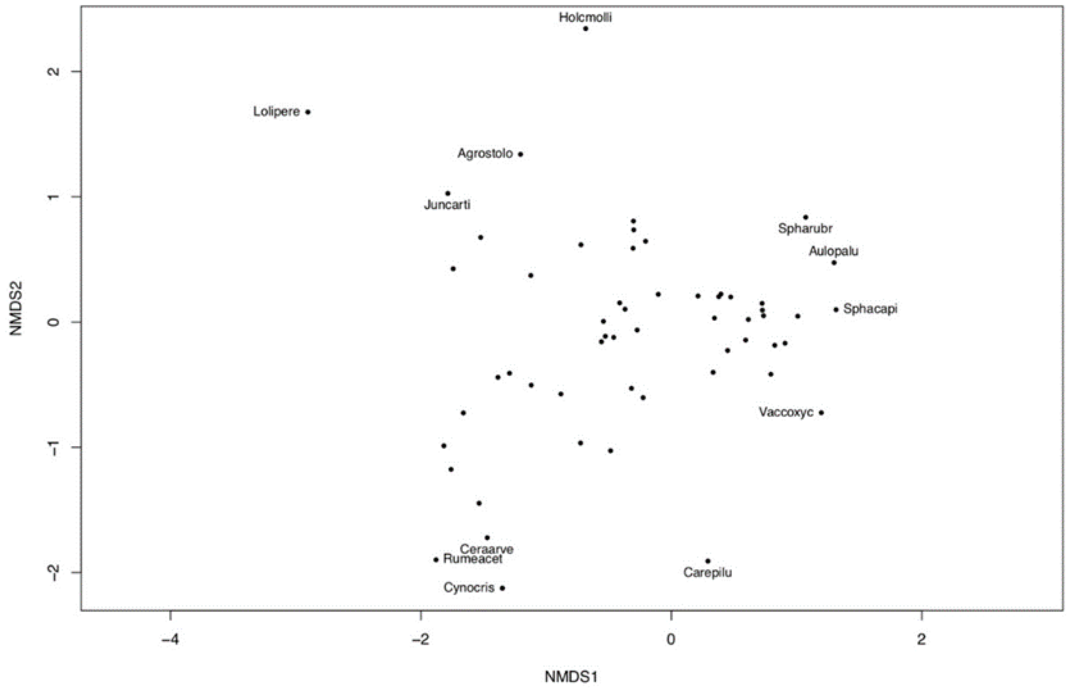


**Fig. S.1c** NMDS ordination species plot for subdominant vegetation found at Ashtrees. Holcmolli = *Holcus lanatus*, Spharubr = *Sphagnum rubra*, Aulopalu = *Aulocomnium palustre*, Sphacapi = *Sphagnum capillofolium*, Vaccoxyc = *Vaccinium oxycoccos*, Carepilu = *Carex pilulifera*, Cynocris = *Cynosurus cristatus*, Rumeacet = *Rumex acetosa*, Ceraarve = *Cirsium arvense*, Juncarti = *Juncus articulatus*, Agrostolo = *Agrostis stolonifera*, Lolipere = *Lolium perenne*

**Table S1. 1** ManyGLM analysis of number of patches created by dominant vegetation and significance of interaction with primary environmental factors. Green = positive correlation, red = negative correlation. (p-values)

| Env. variable | Overall | Callvulg | Nardstri | Molicaer | Eriovagi | Juncsqua | Junceffu | Carenigr | Galisaxa | Poteerec | Descflex | Vaccmyrt |
|---|---|---|---|---|---|---|---|---|---|---|---|---|
| Soil pH | 0.001 | 0.424 | 0.999 | 0.421 | 0.591 | 0.941 | 0.999 | 0.993 | 0.967 | 0.999 | 0.135 | 0.889 |
| Slope | 0.001 | 1.000 | 0.044 | 1.000 | 0.003 | 0.991 | 1.000 | 1.000 | 0.112 | 0.999 | 1.000 | 1.000 |
| % water | 0.001 | 0.999 | 0.998 | 1.000 | 0.001 | 0.999 | 0.998 | 1.000 | 1.000 | 0.999 | 1.000 | 0.999 |
| Altitude | 0.001 | 0.966 | 1.000 | 1.000 | 1.000 | 1.000 | 0.075 | 1.000 | 0.985 | 0.899 | 1.000 | 1.000 |

**Table S1. 2** ManyGLM analysis of area occupied by dominant vegetation and significance of interaction with primary environmental factors. Green = positive correlation, red = negative correlation. (p-values)

| Env. variable | Overall | Callvulg | Nardstri | Molicaer | Eriovagi | Juncsqua | Junceffu | Carenigr | Galisaxa | Poteerec | Descflex | Vaccmyrt |
|---|---|---|---|---|---|---|---|---|---|---|---|---|
| Soil pH | 0.001 | 0.880 | 0.969 | 0.281 | 0.880 | 0.969 | 0.997 | 0.997 | 0.988 | 1.000 | 0.443 | 0.816 |
| Slope | 0.069 | 0.987 | 0.981 | 1.000 | 0.998 | 0.997 | 1.00 | 1.000 | 0.999 | 0.998 | 0.998 | 1.000 |
| % water | 0.029 | 0.934 | 0.960 | 1.000 | 0.039 | 0.932 | 1.00 | 1.000 | 1.000 | 0.977 | 0.039 | 1.000 |
| Altitude | 0.053 | 0.703 | 0.998 | 1.000 | 0.998 | 1.000 | 0.122 | 1.000 | 1.000 | 0.998 | 0.988 | 1.000 |

**Table S1. 3** ManyGLM analysis of shape index for dominant vegetation and significance of interaction with primary environmental factors. Green = positive correlation, red = negative correlation. (p-values)

| Env. variable | Overall | Callvulg | Nardstri | Molicaer | Eriovagi | Juncsqua | Junceffu | Carenigr | Galisaxa | Poteerec | Descflex | Vaccmyrt |
|---|---|---|---|---|---|---|---|---|---|---|---|---|
| Soil pH | 0.001 | 1.000 | 1.000 | 1.000 | 0.842 | 1.000 | 0.974 | 0.992 | 0.720 | 1.000 | 0.072 | 0.914 |
| Slope | 0.001 | 1.000 | 0.417 | 1.000 | 0.022 | 0.995 | 1.000 | 1.000 | 0.736 | 1.000 | 1.000 | 1.000 |
| % water | 0.001 | 1.000 | 1.000 | 1.000 | 0.001 | 0.933 | 1.000 | 0.563 | 0.939 | 1.000 | 0.997 | 0.935 |
| Altitude | 0.048 | 1.000 | 1.000 | 1.000 | 1.000 | 1.000 | 0.016 | 1.000 | 1.000 | 0.929 | 1.000 | 1.000 |

**Table S1. 4** ManyGLM analysis of number of patches created by subdominant vegetation and significance of interaction with primary environmental factors. Green = positive correlation, red = negative correlation. (p-values)

| Env. variable | Overall | Callvulg | Nardstri | Molicaer | Eriovagi | Juncsqua | Junceffu | Carenigr | Galisaxa | Poteerec | Descflex | Vaccmyrt |
|---|---|---|---|---|---|---|---|---|---|---|---|---|
| Soil pH | 0.001 | 0.258 | 1.000 | 0.043 | 0.854 | 1.000 | 1.000 | 1.000 | 0.996 | 1.000 | 0.003 | 0.720 |
| Slope | 0.019 | 1.000 | 0.842 | 1.000 | 0.891 | 0.976 | 1.000 | 1.000 | 0.805 | 1.000 | 0.998 | 1.000 |
| % water | 0.002 | 0.526 | 0.672 | 1.000 | 0.005 | 0.999 | 0.969 | 1.000 | 0.808 | 1.000 | 1.000 | 0.977 |
| Altitude | 0.011 | 0.423 | 1.000 | 1.000 | 1.000 | 1.000 | 0.116 | 1.000 | 1.000 | 0.997 | 0.999 | 1.000 |

**Table S1. 5** ManyGLM analysis of <u>area</u> occupied by <u>subdominant</u> vegetation and significance of interaction with primary environmental factors. Green = positive correlation, red = negative correlation. (p-values)

| Env. variable | Overall | Callvulg | Nardstri | Molicaer | Eriovagi | Juncsqua | Junceffu | Carenigr | Galisaxa | Poteerec | Descflex | Vaccmyrt |
|---|---|---|---|---|---|---|---|---|---|---|---|---|
| Soil pH | 0.001 | 0.178 | 0.999 | 0.574 | 0.919 | 0.984 | 1.000 | 0.993 | 0.991 | 1.000 | 0.014 | 0.990 |
| Slope | 0.036 | 1.000 | 0.957 | 1.000 | 0.975 | 0.996 | 1.000 | 0.858 | 1.000 | 1.000 | 1.000 | 1.000 |
| % water | 0.033 | 0.996 | 0.986 | 0.972 | 0.064 | 0.997 | 1.000 | 1.000 | 1.000 | 1.000 | 1.000 | 0.993 |
| Altitude | 0.066 | 0.361 | 0.996 | 0.999 | 1.000 | 0.999 | 1.000 | 0.668 | 1.000 | 0.955 | 1.000 | 1.000 |

**Table 1. 6** ManyGLM analysis of <u>shape index</u> for <u>subdominant</u> vegetation and significance of interaction with primary environmental factors. Green = positive correlation, red = negative correlation. (p-values)

| Env. variable | Overall | Callvulg | Nardstri | Molicaer | Eriovagi | Juncsqua | Junceffu | Carenigr | Galisaxa | Poteerec | Descflex | Vaccmyrt |
|---|---|---|---|---|---|---|---|---|---|---|---|---|
| Soil pH | 0.001 | 0.414 | 0.991 | 1.000 | 0.878 | 1.000 | 1.000 | 1.000 | 0.873 | 1.000 | 1.000 | 1.000 |
| Slope | 0.001 | 1.000 | 0.102 | 1.000 | 0.102 | 0.950 | 1.000 | 1.000 | 0.164 | 1.000 | 1.000 | 0.983 |
| % water | 0.001 | 0.256 | 0.033 | 1.000 | 0.001 | 0.978 | 0.506 | 1.000 | 0.881 | 1.000 | 1.000 | 0.986 |
| Altitude | 0.001 | 0.973 | 0.983 | 1.000 | 1.000 | 1.000 | 0.256 | 1.000 | 1.000 | 0.997 | 1.000 | 1.000 |

**Table S1. 7** ManyGLM analysis of <u>number of patches</u> for <u>dominant</u> vegetation and correlation with length of sheep track and distance to sheep track within a 10m, 25m and 35m buffer and distance to nearest ditch. ST = Sheep Track. Green = positive correlation, red = negative correlation. (p-values)

| No. of Patches | overall | Callvulg | Eriovagi | Junceffu | Juncsqua | Molicaer | Nardstri | Carenigr | Descflex | Galisaxa | Poteerec | Vaccmyrt |
|---|---|---|---|---|---|---|---|---|---|---|---|---|
| ST length (10m buffer) | 0.720 | 0.998 | 1.000 | 0.998 | 1.000 | 1.000 | 1.000 | 0.998 | 1.000 | 0.987 | 1.000 | 0.997 |
| ST distance (10m buffer) | 0.024 | 1.000 | 1.000 | 0.959 | 0.998 | 1.000 | 1.000 | 0.998 | 1.000 | 1.000 | 0.910 | 1.000 |
| ST length (25m buffer) | 0.06 | 1.000 | 1.000 | 0.910 | 1.000 | 0.982 | 1.000 | 1.000 | 1.000 | 0.998 | 0.781 | 1.000 |
| ST distance (25m buffer) | 0.110 | 1.000 | 1.000 | 1.000 | 1.000 | 0.740 | 0.985 | 0.999 | 1.000 | 0.953 | 1.000 | 0.997 |
| ST length (35m buffer) | 0.200 | 1.000 | 1.000 | 1.000 | 1.000 | 1.000 | 1.000 | 1.000 | 1.000 | 1.000 | 1.000 | 1.000 |
| ST distance (35m buffer) | 0.071 | 1.000 | 1.000 | 1.000 | 1.000 | 1.000 | 1.000 | 1.000 | 1.000 | 1.000 | 0.957 | 1.000 |
| Ditch distance | 0.066 | 1.000 | 1.000 | 1.000 | 1.000 | 1.000 | 1.000 | 1.000 | 1.000 | 0.988 | 0.849 | 0.999 |

**Table S1. 8** ManyGLM analysis of area of patches for dominant vegetation and correlation with length of sheep track and distance to sheep track within a 10m, 25m and 35m buffer and distance to nearest ditch. ST = Sheep Track. Green = positive correlation, red = negative correlation. (p-values)

| Area | overall | Callvulg | Eriovagi | Junceffu | Juncsqua | Molicaer | Nardstri | Carenigr | Descflex | Galisaxa | Poteerec | Vaccmyrt |
|---|---|---|---|---|---|---|---|---|---|---|---|---|
| ST length (10m buffer) | 0.430 | 1.000 | 0.979 | 0.996 | 1.000 | 0.892 | 1.000 | 1.000 | 1.000 | 0.966 | 1.000 | 0.998 |
| ST distance (10m buffer) | 0.085 | 1.000 | 1.000 | 0.944 | 0.998 | 1.000 | 1.000 | 0.962 | 1.000 | 1.000 | 0.998 | 1.000 |
| ST length (25m buffer) | 0.544 | 1.000 | 1.000 | 1.000 | 1.000 | 0.651 | 1.000 | 1.000 | 1.000 | 1.000 | 1.000 | 1.000 |
| ST distance (25m buffer) | 0.875 | 1.000 | 1.000 | 1.000 | 1.000 | 1.000 | 1.000 | 1.000 | 1.000 | 1.000 | 1.000 | 1.000 |
| ST length (35m buffer) | 0.169 | 1.000 | 1.000 | 1.000 | 1.000 | 1.000 | 1.000 | 1.000 | 1.000 | 1.000 | 1.000 | 1.000 |
| ST distance (35m buffer) | 0.023 | 1.000 | 1.000 | 1.000 | 1.000 | 1.000 | 1.000 | 0.961 | 1.000 | 1.000 | 0.866 | 0.996 |
| Ditch distance | 0.167 | 1.000 | 0.941 | 1.000 | 1.000 | 1.000 | 1.000 | 1.000 | 1.000 | 1.000 | 0.963 | 1.000 |

**Table S1. 9** ManyGLM analysis of shape index of patches for dominant vegetation and correlation with length of sheep track and distance to sheep track within a 10m, 25m and 35m buffer and distance to nearest ditch. ST = Sheep Track. Green = positive correlation, red = negative correlation. (p-values)

| Shape | overall | Callvulg | Eriovagi | Junceffu | Juncsqua | Molicaer | Nardstri | Carenigr | Descflex | Galisaxa | Poteerec | Vaccmyrt |
|---|---|---|---|---|---|---|---|---|---|---|---|---|
| ST length (10m buffer) | 0.533 | 1.000 | 0.976 | 1.000 | 1.000 | 0.976 | 0.624 | 0.998 | 1.000 | 1.000 | 1.000 | 1.000 |
| ST distance (10m buffer) | 0.478 | 0.999 | 1.000 | 1.000 | 1.000 | 0.579 | 0.997 | 0.999 | 1.000 | 0.992 | 1.000 | 0.999 |
| ST length (25m buffer) | 0.055 | 1.000 | 1.000 | 1.000 | 0.999 | 1.000 | 1.000 | 0.999 | 1.000 | 1.000 | 0.784 | 1.000 |
| ST distance distance (25m buffer) | 0.289 | 1.000 | 1.000 | 1.000 | 1.000 | 1.000 | 1.000 | 1.000 | 1.000 | 1.000 | 1.000 | 1.000 |
| ST length (35m buffer) | 0.097 | 1.000 | 1.000 | 1.000 | 1.000 | 1.000 | 1.000 | 1.000 | 1.000 | 1.000 | 1.000 | 1.000 |
| ST distance (35m buffer) | 0.059 | 0.979 | 1.000 | 1.000 | 1.000 | 1.000 | 0.972 | 1.000 | 1.000 | 1.000 | 0.996 | 1.000 |
| Ditch distance | 0.023 | 1.000 | 1.000 | 1.000 | 1.000 | 1.000 | 1.000 | 1.000 | 1.000 | 1.000 | 1.000 | 0.992 |

**Table S1. 10** ManyGLM analysis of number of patches for subdominant vegetation and correlation with length of sheep track and distance to sheep track within a 10m, 25m and 35m buffer and distance to nearest ditch. ST = Sheep Track. Green = positive correlation, red = negative correlation. (p-values)

| No. of Patches | overall | Callvulg | Eriovagi | Junceffu | Juncsqua | Molicaer | Nardstri | Carenigr | Descflex | Galisaxa | Poteerec | Vaccmyrt |
|---|---|---|---|---|---|---|---|---|---|---|---|---|
| ST length (10m buffer) | 0.436 | 0.955 | 0.390 | 1.000 | 1.000 | 1.000 | 1.000 | 1.000 | 1.000 | 1.000 | 1.000 | 0.997 |
| ST distance (10m buffer) | 0.048 | 0.630 | 1.000 | 1.000 | 1.000 | 1.000 | 1.000 | 0.930 | 1.000 | 1.000 | 1.000 | 1.000 |
| ST length (25m buffer) | 0.281 | 1.000 | 0.997 | 0.979 | 1.000 | 1.000 | 1.000 | 0.795 | 1.000 | 1.000 | 0.990 | 1.000 |
| ST distance distance (25m buffer) | 0.466 | 1.000 | 1.000 | 1.000 | 1.000 | 1.000 | 1.000 | 1.000 | 1.000 | 1.000 | 1.000 | 1.000 |
| ST length (35m buffer) | 0.206 | 1.000 | 1.000 | 1.000 | 0.988 | 1.000 | 1.000 | 1.000 | 1.000 | 1.000 | 1.000 | 1.000 |
| ST distance (35m buffer) | 0.205 | 0.986 | 1.000 | 1.000 | 1.000 | 1.000 | 1.000 | 1.000 | 1.000 | 1.000 | 1.000 | 1.000 |
| Ditch distance | 0.063 | 1.000 | 1.000 | 0.910 | 1.000 | 1.000 | 1.000 | 1.000 | 1.000 | 0.997 | 0.882 | 1.000 |

**Table S1. 11** ManyGLM analysis of area of patches for subdominant vegetation and correlation with length of sheep track and distance to sheep track within a 10m, 25m and 35m buffer and distance to nearest ditch. ST = Sheep Track. Green = positive correlation, red = negative correlation. (p-values)

| Area | overall | Callvulg | Eriovagi | Junceffu | Juncsqua | Molicaer | Nardstri | Carenigr | Descflex | Galisaxa | Poteerec | Vaccmyrt |
|---|---|---|---|---|---|---|---|---|---|---|---|---|
| ST length (10m buffer) | 0.362 | 0.953 | 0.394 | 1.000 | 1.000 | 1.000 | 1.000 | 1.000 | 1.000 | 0.678 | 1.000 | 1.000 |
| ST distance (10m buffer) | 0.052 | 0.627 | 1.000 | 1.000 | 1.000 | 1.000 | 1.000 | 0.952 | 1.000 | 1.000 | 1.000 | 0.998 |
| ST length (25m buffer) | 0.346 | 1.000 | 1.000 | 0.984 | 1.000 | 1.000 | 1.000 | 0.762 | 1.000 | 1.000 | 0.966 | 1.000 |
| ST distance distance (25m buffer) | 0.400 | 1.000 | 1.000 | 1.000 | 1.000 | 1.000 | 1.000 | 1.000 | 1.000 | 1.000 | 1.000 | 1.000 |
| ST length (35m buffer) | 0.137 | 1.000 | 1.000 | 1.000 | 0.995 | 1.000 | 1.000 | 1.000 | 1.000 | 1.000 | 1.000 | 0.997 |
| ST distance (35m buffer) | 0.196 | 1.000 | 1.000 | 1.000 | 1.000 | 1.000 | 1.000 | 1.000 | 1.000 | 1.000 | 1.000 | 1.000 |
| Ditch distance | 0.108 | 1.000 | 1.000 | 1.000 | 1.000 | 1.000 | 1.000 | 1.000 | 1.000 | 1.000 | 0.937 | 1.000 |

**Table S1. 12** ManyGLM analysis of shape index of patches for subdominant vegetation and correlation with length of sheep track and distance to sheep track within a 10m, 25m and 35m buffer and distance to nearest ditch. ST = Sheep Track. Green = positive correlation, red = negative correlation. (p-values).

| Shape | overall | Callvulg | Eriovagi | Junceffu | Juncsqua | Molicaer | Nardstri | Carenigr | Descflex | Galisaxa | Poteerec | Vaccmyrt |
|---|---|---|---|---|---|---|---|---|---|---|---|---|
| ST length (10m buffer) | 0.362 | 0.953 | 0.394 | 1.000 | 1.000 | 1.000 | 0.981 | 1.000 | 1.000 | 0.995 | 1.000 | 1.000 |
| ST distance (10m buffer) | 0.052 | 0.627 | 1.000 | 1.000 | 1.000 | 1.000 | 1.000 | 0.952 | 1.000 | 1.000 | 1.000 | 0.998 |
| ST length (25m buffer) | 0.346 | 1.000 | 1.000 | 0.984 | 1.000 | 1.000 | 1.000 | 0.762 | 1.000 | 1.000 | 1.000 | 1.000 |
| ST distance distance (25m buffer) | 0.420 | 1.000 | 1.000 | 1.000 | 1.000 | 1.000 | 1.000 | 1.000 | 1.000 | 1.000 | 0.966 | 1.000 |
| ST length (35m buffer) | 0.085 | 1.000 | 1.000 | 1.000 | 1.000 | 1.000 | 1.000 | 1.000 | 1.000 | 1.000 | 1.000 | 0.984 |
| ST distance (35m buffer) | 0.202 | 1.000 | 1.000 | 1.000 | 1.000 | 1.000 | 1.000 | 1.000 | 1.000 | 0.984 | 1.000 | 1.000 |
| Ditch distance | 0.086 | 1.000 | 1.000 | 0.989 | 1.000 | 1.000 | 1.000 | 1.000 | 1.000 | 1.000 | 0.881 | 1.000 |
